# The activity of yeast Apn2 AP endonuclease at uracil-derived AP sites is dependent on the major carbon source

**DOI:** 10.1101/2020.09.01.278135

**Authors:** Kasey Stokdyk, Alexandra Berroyer, Zacharia A. Grami, Nayun Kim

## Abstract

Yeast Apn2 is an AP endonuclease and DNA 3’-diesterase that belongs to the Exo III family with homology to the *E. coli* exonuclease III, *Schizosaccharomyces pombe* eth1, and human AP endonucleases APEX1 and APEX2. In the absence of Apn1, the major AP endonuclease in yeast, Apn2 can cleave the DNA backbone at an AP lesion initiating the base excision repair pathway. In order to study the role and relative contribution of Apn2, we took advantage of a reporter system that was previously used to delineate how uracil-derived AP sites are repaired. At this reporter, disruption of the Apn1-initiated base excision repair pathway led to a significant elevation of A:T to C:G transversions. Here we show that such highly elevated A:T to C:G transversion mutations associated with uracil residues in DNA are abolished when *apn1*Δ yeast cells are grown in glucose as the primary carbon source. We also show that the disruption of Apn2, either by the complete gene deletion or by the mutation of a catalytic residue, results in a similarly reduced rate of the uracil-associated mutations. Overall, our results indicate that Apn2 activity is regulated by the glucose repression pathway in yeast.

## INTRODUCTION

Apurinic/apyrimidic (AP) sites are one of the most common types of endogenous DNA damage and are generated by the cleavage of the N-glycosidic bond between the base and the deoxyribose moiety in DNA by DNA glycosylases (Boiteux and Jinks-Robertson 2013; Wallace 2014). Substrates of DNA N-glycosylases include oxidation-damaged bases, uracil, and spontaneous or enzymatic deamination products. After cleavage of the glycosidic bond, sugar phosphate backbone of the DNA strand adjacent to the AP site is cleaved by AP endonucleases to generate a DNA nick to initiate Base Excision Repair (BER). Yeast contains two structurally unrelated proteins capable of incision at the AP sites, Apn1 and Apn2 (Daley et al. 2010). Apn1 belongs to the Endo IV family and comprises ~97% of both AP endonuclease and DNA 3’ diesterase activity in yeast cell extracts (Popoff et al. 1990). Apn2, which also contains AP endonuclease and DNA 3’-diesterase activity, belongs to the Exo III family with homology to the *E. coli* exonuclease III, *Schizosaccharomyces pombe* eth1, and human AP endonucleases APEX1 and APEX2 (Bennett 1999; Johnson et al. 1998; Unk et al. 2000). Endo IV family AP endonucleases are not found in most metazoan species including insects and mammals (Daley et al. 2010).

Uracil is one of the most frequent types of endogenous DNA modifications that are normally repaired by BER (Collura et al. 2012; Guillet and Boiteux 2003). The spontaneous appearance of uracil in DNA can be attributed to the eukaryotic replicative DNA polymerases that cannot distinguish between a uracil and a thymine base (Bessman et al. 1958; Warner et al. 1981). Depending on the cellular dUTP concentration, uracil is readily incorporated in place of thymine during replication and repair, resulting in a stable U:A base pair. An abasic site generated through the removal of uracil by a uracil DNA glycosylase (Ung1 in yeast; UDG in human) are either repaired in error-free manner by BER machinery or result in further mutagenic event mediated by the error-prone translesion synthesis DNA polymerases (Boiteux and Jinks-Robertson 2013). When left unrepaired, the nick left by AP endonucleases can result in DNA strand breaks leading to more detrimental genome instability events (Luke et al. 2010). Significantly, genome instability induced by uracil in DNA is the principal cause of the cytotoxicity of two major classes of anticancer drugs. The fluoropyrimidines, 5-fluorouracil (5-FU) and capecitabine are metabolized into 5-flurorodeoxyuridylate, which binds the essential enzyme thymidylate synthase (TS) to form an inhibitory complex (Longley et al. 2003). Antifolates (e.g. methotrexate) disrupt TS activity by inhibiting the enzyme dihydrofolate reductase (DHFR) and thereby interfering with the production of required co-factor tetrahydrofolate (Hagner and Joerger 2010). Because TS is the sole enzyme catalyzing the production of thymidylate (dTTP) from uracil (dUTP), in cells treated with fluoropyrimidines or antifolates, relatively lower dTTP concentration forces elevated incorporation of dUTP during replication and subsequent genome instability.

We have recently demonstrated a correlation between the elevated levels of uracil-derived mutations at highly transcribed genes and the higher densities of uracil residues in genomic DNA (Kim et al. 2007; Kim and Jinks-Robertson 2009; Kim and Jinks-Robertson 2010; Owiti et al. 2018). To study how uracil in DNA are incorporated and repaired, we constructed several mutagenesis reporters designed to study the rate and the spectrum of mutations associated with active transcription in the yeast genome. One of these, the *pTET-lys2-TAA* reporter, contains an in-frame insertion of a TAA stop codon within the coding region of *LYS2* gene. Thus, yeast cells harboring this allele are lysine auxotrophs (Lys^-^) and reverts to lysine prototrophs (Lys^+^) through mutations at the TAA stop codon allowing translation read-through. It was shown that a great majority of Lys^+^ reversion mutations occurring at the *pTET-lys2-TAA* are due to the unrepaired AP lesions arising from the excision of uracil in DNA (Kim and Jinks-Robertson 2009; Kim and Jinks-Robertson 2010). When the *pTET-lys2-TAA* was actively transcribed, there was a sharp elevation in A to C or T to G transversion mutations. These mutations were dependent on the activity of translesion-synthesis (TLS) polymerases Pol zeta and Rev3, which together insert a C nucleotide opposite the AP sites (Kim et al. 2011b). And, these mutations were significantly reduced by the overexpression of dUTP pyrophosphatase Dut1 and mostly eliminated by the disruption of the uracil-specific DNA glycosylase Ung1 (Kim and Jinks-Robertson 2009; Kim and Jinks-Robertson 2010). Because Dut1 performs an important function of regulating the cellular concentration of dUTP by converting dUTP to dUMP and inorganic pyrophosphate, its overexpression leads to lower dUTP pool available for incorporation into the genome (Guillet et al. 2006). And in the absence of Ung1, uracil remains in the genomic DNA without triggering base excision repair (BER) or mutagenic TLS. Overall, these data indicate that the transcription-associated A:T to C:G transversions at the *pTET-lys2-TAA* reporter result from the TLS-mediated bypass of AP lesions produced through uracil-excision by Ung1. The uracil-derived A:T to C:G mutations are highly elevated in strains deficient for Apn1 protein.

Here we used the previously-established genetic tools to study the contribution of Apn2 AP endonuclease in the repair of uracil-derived AP lesions in yeast. Our data show that Apn2 enhances the mutations associated with uracil-derived AP sites in yeast in a manner dependent on the available carbon source. This surprising result calls for further study into how the activity of Apn2 is regulated by the presence of different carbon sources and by glucose in particular.

## MATERIALS AND METHODS

### Yeast strains and plasmids

Yeast strains used for the mutation and recombination assays were derived from YPH45 (*MATa, ura3-52 ade2-101 trp1Δ1*). Construction of strains containing the *his4Δ::pTET-lys2-TAA* allele was previously described (Kim and Jinks-Robertson 2010). See the strain list in Table S1 for more information. Gene deletions were carried out through one-step allele replacement by amplification of loxP-flanked marker cassettes (Gueldener et al. 2002). The entire *APN2* ORF containing the E59A mutation was synthesized by the GeneArt Gene Synthesis service (ThermoFisher Scientific) and cloned into a pGPD2 vector (Addgene #43972) to construct pNK231. YNK650 containing *apn2 E59A* allele was constructed through pop-in/pop-out method using BseRI-digested pNK231. For overexpression of *APN2* gene, the *APN2* ORF was amplified from the yeast genomic DNA and cloned between XbaI and XhoI sites of pGPD2 vector to construct pNK229.

### Mutation/Recombination Rates and Spectra

Mutation and recombination rates were determined using the method of the median, and 95% confidence intervals were calculated as previously described (Spell and Jinks-Robertson 2004). Two rates are considered significantly different when the confidence intervals do not overlap. Each rate was based on data obtained from 12-48 independent cultures and at least two independently-derived isolates. 1mL yeast extract-peptone (YEP) medium supplemented with 2% glycerol and 2% ethanol (YEPGE) or with 2% dextrose (YEPD) was inoculated with 250,000 cells from an overnight culture grown in the same medium. Following growth at 30°C for 2 days in YEPD or for 3 days in YEPGE, cells were washed with water and the appropriate dilutions were plated either on synthetic, lysine-deficient medium containing 2% glucose (SCD-Lys) to select for Lys+ revertants or on SCD-Leu medium to determine the total number of cells in each culture. *CAN1* forward mutation rates were determined by plating cells on SCD-Arg medium supplemented with 60μg/mL L-canavanine sulfate (SCD-Arg+Can; Sigma). For complementation studies in Figure 2B, YNK103 (*apn1Δ:: loxP apn2*Δ*::loxP-TRP1-loxP*) was transformed with either the vector (pGPD2; Addgene #43972) or pNK229 (pGPD-APN2) and cultured in synthetic, uracil-deficient medium (SC-URA) containing either 2% glucose or 2% glycerol + 2% ethanol at 30°C for 3 days. Then, cells were washed with water and the appropriate dilutions were plated either on synthetic, lysine, uracil-deficient medium containing 2% glucose (SCD-Lys-URA) to select for Lys+ revertants or on SCD-URA medium to determine the total number of cells in each culture.

To determine mutation spectra, individual colonies were used to inoculate 0.3ml liquid YEPGE or YEPD media. After 2 or 3 days of growth at 30°C, an appropriate fraction of each culture was plated on SCD-Lys. A single Lys+ revertant from each culture was purified on YEPD plates, and genomic DNA was prepared using a 96-well format in microtiter plates. The *lys2-TAA* reversion window was amplified using primers 5’-AGCTCGATGTGCCTCATGATAG-3’ and 5’-CATCACACATACCATCAAATCC-3’ and the PCR product was sent to Eurofins Genomics (Louisville, KY) for sequencing using primer 5’-TAGAGTAACCGGTGACGATG-3’. The rates of A>C and T>G were calculated by multiplying the proportion of the events by the total Lys+ mutation rate.

### qRT-PCR

Total RNA was extracted using the standard hot acid phenol method and treated with DNase 1 (New England Biolabs). qRT-PCR was performed using amfiRivert cDNA synthesis Platinum Master Mix from Gendepot, SensiFAST SYBR No-ROX kit from Bioline, and a Biorad CFX Connect instrument. The qRT-PCR conditions were as follows: 45°C for 10 mins and 95°C for 2 mins followed by 40 cycles of 95°C for 5 sec, 60°C for 10 sec and 72°C for 5 sec. *ALG9* gene was chosen as the reference gene (Teste et al. 2009). The primers used for amplification were APN2F: 5’-TTCGAATTATGGCTCACGGA-3’, APN2R: 5’-ACCAGGTTCAATTCTGTCGT-3’, ALG9F: 5’-CACGGATAGTGGCTTTGGTGAACAATTAC-3’, and ALG9R: 5’-TATGATTATCTGGCAGCAGGAAAGAACTTGGG-3’. Relative RNA levels were determined by *ΔΔCq* analysis as previously described (Livak and Schmittgen 2001).

### Identification of putative SUMO- or Ub-modification sites

For the identification of putative sumoylation sites and SUMO-interacation sites within Apn2 protein, we used the GPS-SUMO webserver (Ren et al. 2009; Zhao et al. 2014). UbPred was used for the search of putative ubiquitination sites (Radivojac et al. 2010).

## RESULTS

### Uracil-derived mutations at a highly transcribed gene are dependent on the growth conditions

In previously reported results, the mutation rate and spectra at the *pTET-lys2-TAA* reporter described in the Introduction were determined with the yeast cells cultured in rich media containing yeast extract and peptone with the addition of 2% glycerol and 2% ethanol as the primary carbon source (YEPGE). Here, we repeated the experiments by culturing the yeast cells with the same rich media containing 2% dextrose (YEPD). Surprisingly, we observed that, in an *apn1*Δ background, the rate of mutations at the *pTET-lys2-TAA* reporter when cultured in YEPD was ~24-fold lower than with YEPGE (Figure 1A).

**Figure 1.**
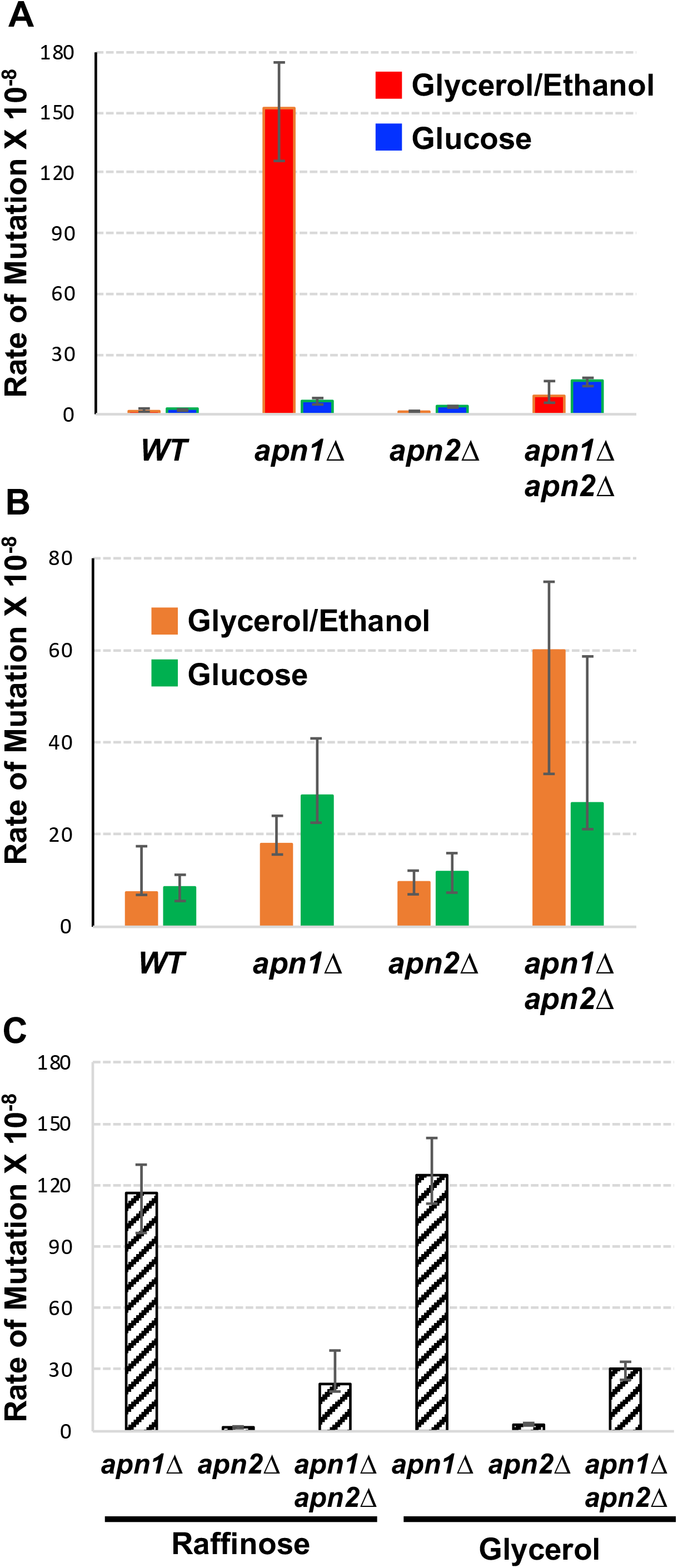
The rates of Lys+ and Can^R^ mutations in various carbon sources. In all graphs, error bars indicate 95% confidence intervals. Two rates are considered significantly different when the error bars do not overlap. **A)** Rates of Lys+ mutations at *pTET-lys2-TAA* in the indicated strain backgrounds. Rates were measured by fluctuation analyses in cells cultured in YEP media supplemented with either 2% Glycerol + 2% ethanol (YEPGE) or 2 % glucose (YEPD). **B)** Rates of canavanine-resistant mutations at *pTET-lys2-TAA* in the indicated strain backgrounds. Rates were measured by fluctuation analyses in cells cultured in YEP media supplemented with either 2% Glycerol + 2% ethanol (YEPGE) or 2 % glucose (YEPD). **C)** Rates of Lys+ mutations at *pTET-lys2-TAA* in the indicated strain backgrounds. Rates were measured by fluctuation analyses in cells cultured in YEP media supplemented with either 2% Glycerol or 2 % raffinose.

**Figure 2.**
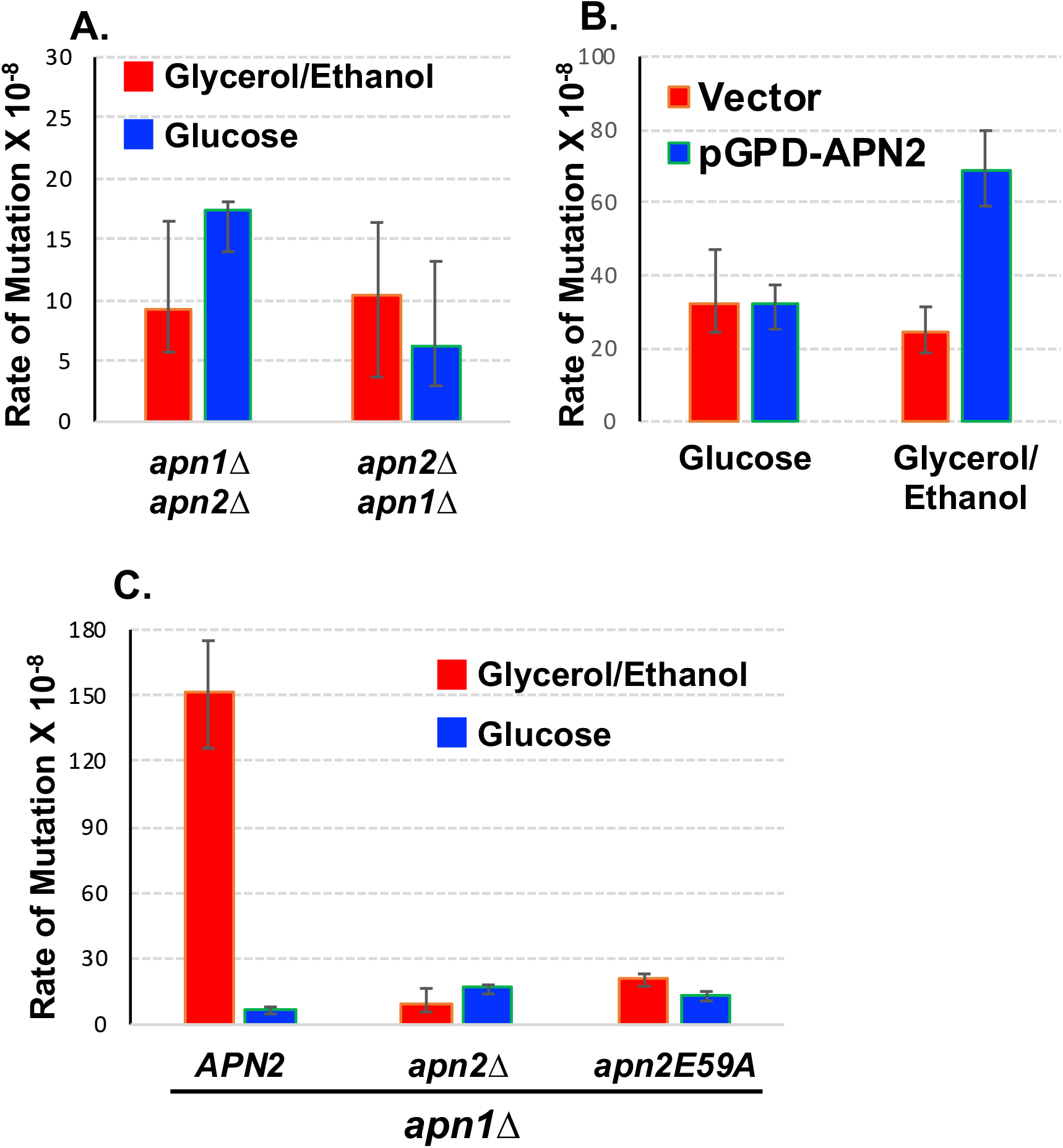
The requirement of Apn2 catalytic activity for the elevated Lys+ mutations in glycerol/ethanol. In all graphs, error bars indicate 95% confidence intervals. Two rates are considered significantly different when the error bars do not overlap. **A)** Rates of Lys+ mutations at *pTET-lys2-TAA* in either *apn1Δ apn2*Δ (YNK103; see Table S1) or *apn2Δ apn1*Δ (YNK219; see Table S1). Rates were measured by fluctuation analyses in cells cultured in YEP media supplemented with either 2% Glycerol + 2% ethanol (YEPGE) or 2 % glucose (YEPD). **B)** Rates of Lys+ mutations at *pTET-lys2-TAA* in YNK103 (*apn1D:: loxP apn2Δ::loxP-TRP1-loxP*) transformed with either the vector (pGPD2; Addgene #43972) or pNK229 (pGPD-APN2). Rates were measured by fluctuation analyses in cells cultured in synthetic, uracil-deficient medium (SC-URA) containing either 2% glucose or 2% glycerol + 2% ethanol as indicated. **C)** Rates of Lys+ mutations at *pTET-lys2-TAA* in YNK98 (*apn1Δ APN2)*, YNK103 (*apn1Δ apn2Δ)*, or YNK658 (*apn1Δ apn2E59A)*. Rates were measured by fluctuation analyses in cells cultured in YEP media supplemented with either 2% Glycerol + 2% ethanol (YEPGE) or 2 % glucose (YEPD).

In a WT background, the deletion of *APN2* gene, which encodes an Exo III family of AP endonuclease, did not result in a significant increase in the mutation at the *pTET-lys2-TAA* reporter either in YEPGE or YEPD media. However, in an *apn1*Δ background, the highly elevated Lys+ reversion mutation rate at the *pTET-lys2-TAA* reporter in cells cultured in YEPGE was reduced by a significant 16-fold reduction upon the deletion of *APN2* (Figure 1A). In addition to the reduction in the overall mutation rate, there was a significant shift in the mutation spectrum upon the deletion of *APN2* in an *apn1*Δ background. The proportion of A:T to C:G transversions, which comprise >90% of the mutations in *apn1*Δ strain, decreased to ~60% of total mutations in the *apn1Δ apn2*Δ strain (Table 1). Upon the deletion of *APN2*, the rates of T>G and A>C mutations in *apn1*Δ cells grown in YEPGE were reduced by 40- and 23-fold, respectively. Interestingly, this effect of *APN2* deletion on the Lys+ reversion mutation rate at the *pTET-lys2-TAA* reporter was entirely dependent on the carbon source available. That is, in YEPD media, the rate of Lys+ reversion mutation was ~3-fold higher in an *apn1Δ apn2*Δ background than in an *apn1*Δ background (Figure 1A). Also in YEPD media, the deletion of *APN2* gene had minimal effect on the mutation spectra in *apn1*Δ cells with the rates of T>G and A>C mutations both increasing by less than 2-folds (Table 1).

**Table 1.**
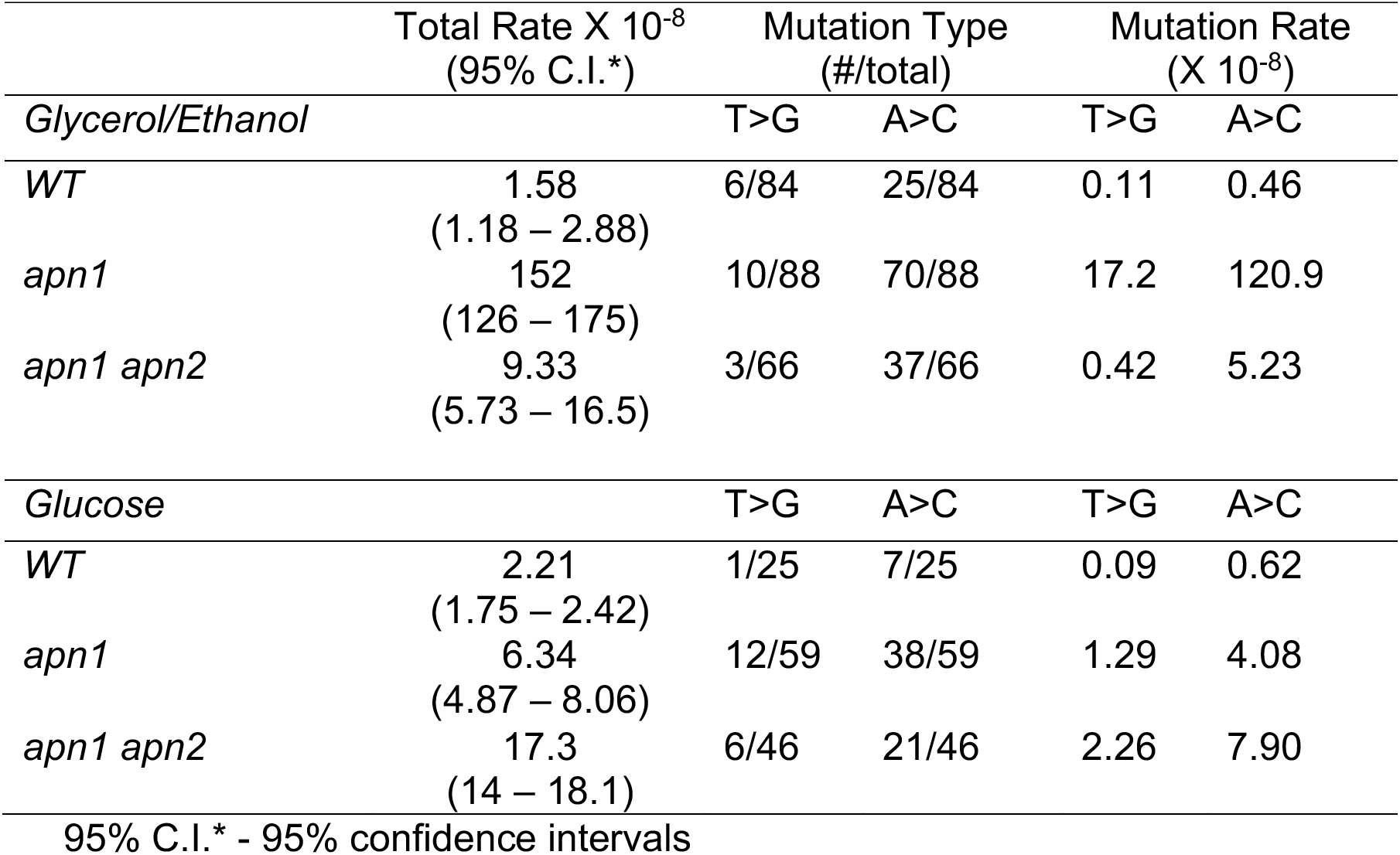
The mutation spectra.

Mutations at the *CAN1* gene that disrupt the function of the encoded arginine transporter protein results in the resistance to the toxic arginine analog canavanine (Whelan et al. 1979). As previously reported (Hanna et al. 2004; Johnson et al. 1998), canavanine-resistance conferring mutations at the *CAN1* gene (Can^R^) are not associated with uracil-derived AP sites. The majority of the base substitutions at *CAN1* genes consists of either C:G to A:T transversions or C:G to T:A transitions. Also as previously reported, we found that the rate of Can^R^ mutation was not affected greatly by the single deletion of *APN1* or *APN2* but synergistically elevated when both are knocked out (Figure 1B). Importantly, no significant difference in the rate of Can^R^ mutation was observed whether the carbon source provided was Glycerol/ethanol or glucose. Thus, the origin of mutation at *CAN1* gene is distinctly different from the uracil-derived mutation occurring at the *pTET-lys2-TAA* reporter under high transcription conditions. Furthermore, while the deletion of *UNG1* gene encoding the uracil DNA glycosylase severely reduces the mutations at the *pTET-lys2-TAA* reporter, the same gene deletion was reported to elevate the rate of spontaneous mutation at *CAN1* gene (Guillet and Boiteux 2003). This further indicates that the elevated mutation rate when cultured in glycerol/ethanol carbon is specific to uracil-derived AP sites.

### Alternative carbon sources do not diminish the mutations at the *pTET-lys2-TAA* reporter

The striking difference in the rate of mutation at the *pTET-lys2-TAA* reporter in YEPD *vs*. YEPGE media led us to investigate the effect of alternate carbon sources. Raffinose is a trisaccharide made up of galactose, glucose, and fructose (Paulo et al. 2015). Raffinose can support growth by fermentation like glucose but favors mitochondrial respiration like glycerol/ethanol (Guaragnella et al. 2013). Global proteomic studies showed that two carbon sources, glucose and raffinose, result in distinctively different protein profiles (Paulo et al. 2015). For cells growing in rich media supplemented with 2% raffinose, the rate of mutation occurring at the *pTET-lys2-TAA* reporter in *apn1*Δ background was statistically the same as that in cells growing in YEPGE (Figure 1C). And similar to the cells in YEPGE, the mutation rate was significantly decreased when *APN2* was further deleted. We also tested whether the ethanol content in the YEPGE media is the decisive factor by repeating the mutation analysis in media containing only 2% glycerol. The rates of mutations at the *pTET-lys2-TAA* reporter were not significantly different from YEPGE in all three backgrounds tested: *apn1*Δ, *apn2*Δ, and *apn1*Δ *apn2*Δ.

### The elevation of uracil-derived A:T to C:G transversions in yeast cells grown in YEPGE requires the catalytic activity of Apn2

In order to first confirm that Apn2 is necessary for the highly elevated mutations at the *pTET-lys2-TAA* reporter in *apn1*Δ background, we independently reconstructed the double knock-out strain by deleting *APN1* gene from the *apn2*Δ strain. The mutation rates at the *pTET-lys2-TAA* reporter in this new strain (*apn2*Δ *apn1*Δ in Figure 2A) were not significantly different from those in the original double knock-out strain (*apn1*Δ *apn2*Δ in Figure 2A), either in YEPD or YEPGE. Similar to the mutation rate in *apn1Δ apn2*Δ cells, the rate of the *pTET-lys2-TAA* mutation in *apn2*Δ *apn1*Δ cells grown in YEPGE was ~14-fold lower and ~9-fold higher when compared to *apn1*Δ and *apn2*Δ single deletion strains, respectively (Figure 2A). Next, we constructed a plasmid transcribing the *APN2* gene from the highly active TDH3 gene promoter which is referred to as GPD promoter (Crook et al. 2011). Overexpression of *APN2* from this plasmid in *apn1*Δ *apn2*Δ strain resulted in a statistically significant, ~3-fold elevation in the mutations at the *pTET-lys2-TAA* reporter in YEPGE but not in YPD media. However, the relative difference in the rate of mutation between *apn1*Δ *apn2*Δ strain with the control vector plasmid and the pGPD-APN2 plasmid is substantially less than the difference between *apn1*Δ and *apn1*Δ *apn2*Δ. This difference is likely an artifact of the overexpression system.

In order to determine whether the effect on the mutation rates at the *pTET-lys2-TAA* reporter requires the catalytic activity of Apn2 protein, we introduced an allelic mutation to change the catalytic glutamate residue E59 of Apn2 to alanine (e.g. *apn2 E59A* allele) (Li et al. 2019). In the strain expressing the catalytically inactive Apn2 E59A mutant protein in the *apn1*Δ background (*apn1*Δ *apn2 E59A*), the mutation rate at the *pTET-lys2-TAA* reporter was significantly lower than that in the *apn1*Δ strain and comparable to that in the *apn1*Δ *apn2*Δ strain (Figure 2B), indicating that the catalytic activity of Apn2 endonuclease is necessary for the elevated uracil-derived A:T to C:G transversions in yeast cells grown in YEPGE. E59A mutation, however, did not affect the mutation rate in YEPD media.

Since Apn2 protein is required for the elevated mutations at the *pTET-lys2-TAA* reporter, the difference in the rate of mutations between YEPD and YEPGE could be due to the level of *APN2* transcription in cells under these different growth conditions. Using qRT-PCR method, we quantified the level of *APN2* transcripts in cell grown in YEPD or YEPGE. As shown in Table 2, either in *WT* or *apn1*Δ background, the difference in the *APN2* gene expression level were not very significant. Although there was a moderately lower Apn2 transcript level in YEPD compared to YEPGE in WT background, there was no significant difference in *apn1*Δ background. This is consistent with the previously reported proteomics study, which showed the difference in the level of Apn2 protein in two different carbon sources was minimal with the level in raffinose being about 73% of the level in glucose (Paulo et al. 2015).

**Table 2.** The rate of *APN2* gene expression.

| Strain Background | Media | Expression Fold Difference Relative to WT cells in YEPGE | Expression Fold Difference Relative to the same background cell in YEPGE |
| --- | --- | --- | --- |
| *WT* | YEPGE | -- | -- |
| *WT* | YPD | 0.58 – 0.64 | 0.58 – 0.64 |
| *apn1Δ* | YEPGE | 1.16 – 1.54 | -- |
| *apn1Δ* | YPD | 1.21 – 1.72 | 0.91 – 1.29 |
| *apn1Δ rfx1Δ* | YEPGE | 4.13 – 10.75 | -- |
| *apn1Δ rfx1Δ* | YPD | 4.73 – 8.72 | 0.71 – 1.31 |

### Stress response genes *YAP1, DUN1*, and *MSN2/MSN4* are not involved in the elevated uracil-derived mutations mediated by Apn2 in YEPGE in an *apn1*Δ background

The expression and activity of many DNA repair proteins are regulated by stress response pathways. We speculated whether a stress response pathway could alter the activity of Apn2, thereby resulting in the highly elevated mutations during growth in YEPGE. We generated yeast strains defective in the stress response genes *YAP1, DUN1*, or *MSN2/MSN4*. Yap1 is a basic leucine zipper transcription factor required for oxidative stress response (Temple et al. 2005) with greater than 500 target genes (Cohen et al. 2002; Venters et al. 2011). *DUN1* encodes a serine/threonine kinase that regulates the DNA damage response (Zhou and Elledge 1993). One of its best characterized targets is the transcriptional repressor Sml1, which in turn regulates the dNTP pools by transcriptional regulation of the ribonucleotide reductase genes. Msn2 and its paralog Msn4 are transcription activator proteins that bind to the stress response elements or STRE (AGGGG or CCCCT) and regulate various stress responses including DNA replication stress and nutrient starvation (Martinez-Pastor et al. 1996; Schmitt and McEntee 1996). For the rate of mutations at the *pTET-lys2-TAA* when in YEPGE, there were no significant changes observed upon the deletion of *YAP1, DUN1*, or *MSN2/4* (Figure 3A, 3C, 3E, and 3G), indicating that none of these three stress response pathways affect the activity of Apn2 in YEPGE. When in YEPD, the rates of mutations at the *pTET-lys2-TAA* were slightly elevated by the deletion of *DUN1* and slightly decreased by the deletion of *MSN2/4* for *apn1*Δ *apn2*Δ background cells (Figure 3D and 3F).

**Figure 3.**
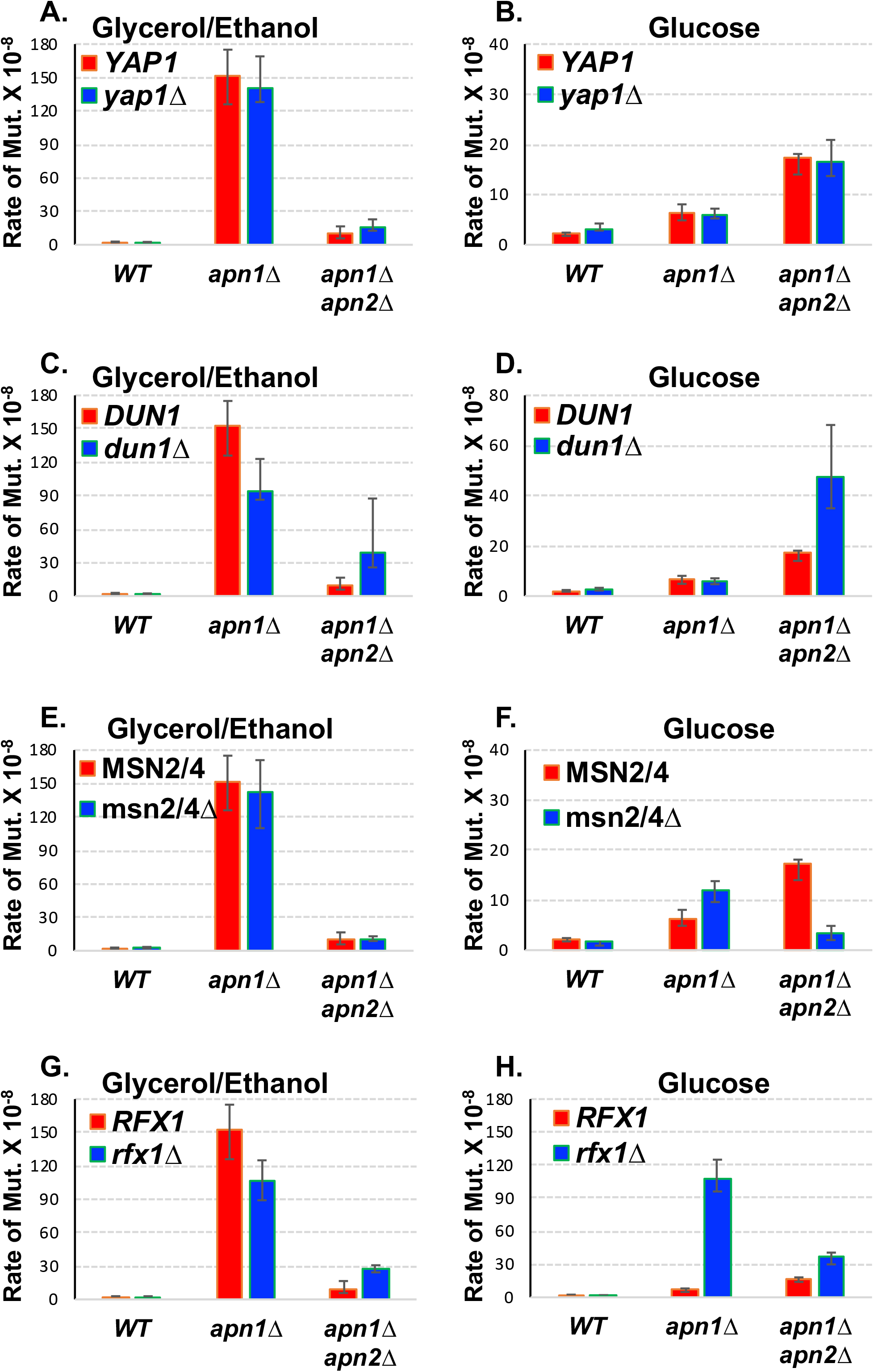
Stress response pathways and the elevated Lys+ mutations in glycerol/ethanol. In all graphs, error bars indicate 95% confidence intervals. Two rates are considered significantly different when the error bars do not overlap. **A)** through **H)** Rates of Lys+ mutations at *pTET-lys2-TAA* in the indicated strain backgrounds. Rates were measured by fluctuation analyses in cells cultured in YEP media supplemented with either 2% Glycerol + 2% ethanol (YEPGE) or 2 % glucose (YEPD) as indicated above each graph. For more information about the strains used, see Table S1.

### The mutation rate at the *pTET-lys2-TAA reporter* becomes elevated upon *RFX1* deletion in YEPD

Rfx1 is another DNA binding transcription factor involved in the DNA damage response. It suppresses the transcription of multiple DNA-damage inducible genes by recruiting general repressor complex Ssn6/Tup1 to the gene promoters (Zhang and Reese 2005). When we deleted *RFX1* gene, the rate of mutations at the *pTET-lys2-TAA* reporter was not significantly changed in *WT, apn1*Δ, or *apn1*Δ *apn2*Δ backgrounds when in YEPGE (Figure 3G). However, in YEPD, the rate of mutations at the *pTET-lys2-TAA* reporter was ~17-fold higher in the *apn1*Δ *rfx1*Δ strain compared to the *apn1*Δ strain (Figure 3H). For *WT* or *apn1*Δ *apn2*Δ background, the deletion of RFX1 gene did not have any effect on the mutation rates at the *pTET-lys2-TAA* reporter. Rfx1 was reported to regulate the level of *APN2* expression during heat-response in a high-throughput analysis (Venters et al. 2011) leading to the speculation that the elevated mutation rate we observed could be due to the alleviation of the transcriptional suppression of *APN2* gene by Rfx1. Using the qRT-PCR approach, we confirmed that the *APN2* expression level is elevated by about 4- to 10-fold in absence of Rfx1 in cells cultured in YEPD (Table 2). However, the *APN2* transcription levels in *rfx1*Δ cells were similarly elevated when cultured in YEPGE. Overall, this data indicates that, when growing in glucose-supplemented media, Rfx1 suppresses the activity of Apn2 required for the elevated Lys+ mutations at the *pTET-lys2-TAA* reporter but not through the direct regulation of the level of *APN2* gene transcription. We speculate that Apn2 activity in glucose-supplemented media is indirectly affected by another factor under the regulation of Rfx1.

## DISCUSSION

Previously, we reported that there is a transcription-associated elevation of A:T to C:G mutations at the *pTET-lys2-TAA* reporter located on the yeast chromosome III (Kim and Jinks-Robertson 2010; Kim et al. 2011b). Multiple lines of evidence indicate that these transversion mutations at this reporter are largely due to AP lesions generated by the excision of uracil base (Kim and Jinks-Robertson 2009; Kim and Jinks-Robertson 2010; Owiti et al. 2018). Briefly, the activity of uracil DNA glycosylase Ung1, which converts uracil in DNA into AP sites, is required for these mutations. Additionally, the level of cellular dUTP is an important determinant in these specific mutations. That is, overexpression of the dUTP-degrading Dut1 enzyme results in a significant decrease in A:T to C:G mutations at the *pTET-lys2-TAA* reporter. These mutations originate from uracil bases incorporated into DNA in place of thymine, which are then excised by Ung1 to generate AP sites. Because the specific A:T to C:G conversions occur when translesion DNA polymerases Pol zeta and Rev1 insert C nucleotides across from unrepaired AP sites during replication, they are highly elevated when BER pathway is disrupted. In yeast, when the major AP endonuclease Apn1 is absent, the repair of AP sites can also be initiated by the AP lyases Ntg1 and Ntg2 (Meadows et al. 2003). It was shown previously that Ntg1/Ntg2-associated BER is preferentially involved in repair of AP sites located on the non-transcribed DNA strand (Kim and Jinks-Robertson 2010). Those AP sites located on the transcribed DNA strand can alternatively be removed by the transcription-coupled nucleotide excision repair pathway (Kim and Jinks-Robertson 2010; Owiti et al. 2017). We have previously reported that A:T to C:G mutations at the *pTET-lys2-TAA* reporter are significantly elevated in *apn1*Δ background and further elevated when the Ntg1/Ntg2 AP lyases are absent or the alternative repair pathway of transcription-coupled nucleotide excision repair (TC-NER) is impaired.

Here we examined whether the other yeast AP endonuclease Apn2 is involved in the repair of AP sites generated by excision of uracil in DNA. Although the deletion of *APN2* gene did not affect the mutations occurring at the *pTET-lys2-TAA* reporter in WT background, a severe decrease in the mutation rate was observed in *apn1*Δ background (Figure 1A). Our data also shows that the catalytic activity of yeast Apn2 is required for the highly elevated A:T to C:G transversions at the actively transcribed *pTET-lys2-TAA* reporter (Figure 2C). Apn1 and Apn2 can each initiate BER by cleavage of the phosphodiester bond adjacent to an AP lesion. Such a redundant function is shown in the rate of Can^R^ mutations as in Figure 1B, which are elevated in *apn1*Δ strain and further elevated in *apn1*Δ *apn2*Δ double mutant strain. On the other hand, the rate of uracil-derived A:T to G:C transversions at the *pTET-lys2-TAA* reporter are significant elevated in *apn1*Δ but reduced in *apn1*Δ *apn2*Δ strain (Figure 1A), indicating the Apn2 activity involved must be a non-canonical function other than the cleavage of the DNA backbone. Recently, there have been reports of such non-canonical function of Apn2. In the absence of RNase H2-mediated ribonucleotide excision repair (RER), Top1-depenent cleavage at ribonucleotide embedded in DNA results in 2’,3’-cyclic phosphate terminated DNA ends, which leads to 2-5 bp deletion mutations at repetitive sequences (Kim et al. 2011a). Apn2 was recently shown to process these blocked 3’ ends produced during the Top1-mediated cleavage of ribonucleotides in DNA (Li et al. 2019). The ribonucleotide-associated 2-5 bp deletions are elevated by >100-fold in absence of Apn2 but not in absence of Apn1. The same catalytic mutation (i.e. E59A) that abolishes A:T to C:G mutations at the *pTET-lys2-TAA* reporter here was shown to also abolish the ribonucleotide-associated small deletions in this study.

Another key finding reported here is that the highly elevated Apn2-dependent A:T to C:G transversions are not observed when the yeast cells are cultured with glucose as the primary carbon source (Figure 1A). While dextrose is present at a sufficient concentration, yeast cells can sustain energy production and growth by fermentation even in the presence of oxygen (Pfeiffer and Morley 2014), while gene involved in the use of other carbon sources are generally suppressed in a process referred to as glucose repression (Kayikci and Nielsen 2015). Deactivation of glucose repression is required for the utilization of non-fermentable carbon sources such as glycerol (Schuller 2003). When growing in raffinose as the major carbon source, ATP production in yeast cells is primarily carried out through mitochondrial respiration and not through fermentation with no evidence of glucose repression (Guaragnella et al. 2013; Randez-Gil et al. 1998). We observed the high A:T to C:G conversion rate in yeast cells growing in media supplemented with 2% glycerol + 2% ethanol, 2% glycerol only, or 2% raffinose only but not with 2% glucose only (Figure 1A and 1C).

The suppression of Apn2 activity in glucose is not at the level of transcriptional regulation. *APN2* transcripts in both WT and *apn1*Δ cells as measured by qRT-PCR are present in similar level whether glucose (YEPD) or glycerol/ethanol (YEPGE) is the major carbon source (Table 2) and thus cannot sufficiently account for the higher mutation rate observed in YEPGE. Either in glucose or glycerol/ethanol, the rate of Lys+ mutations at the *pTET-lys2-TAA* reporter was not significantly affected by disruption of the Yap1, Dun1, or Msn2/4 stress response factors (Figure 3A – 3F). When *RFX1* gene was deleted in *apn1*Δ background, the rate of Lys+ mutations at the *pTET-lys2-TAA* reporter in glucose was highly elevated to be similar to the rate of mutations in glycerol/ethanol (Figure 3G and 3H). However, the *APN2* gene expression is elevated by 4- to 10-fold both in YEPD and YEPGE by *RFX1* gene deletion (Table 2) even though there was no significant increase in the rate of mutation in an *apn1*Δ *rfx1*Δ strain compared to an *apn1*Δ strain in glycerol/ethanol (Figure 3G). This result suggests that the transcriptional regulation of *APN2* does not sufficiently account for its non-canonical activity leading to higher uracil-associated mutations.

One possible way by which Apn2 activity can be regulated is through posttranslation modification. Human AP endonuclease APEX1, which is a class II endonuclease like yeast Apn2, is reported to be regulated by various types of posttranslational modifications (reviewed in (Busso et al. 2010)). For example, the cellular localization of hAPE1 is regulated by ubiquitination of the protein (Busso et al. 2009; Busso et al. 2010). Several putative ubiquitination sites are present in yeast Apn2 according to UbPred web tool (Radivojac et al. 2010) (Figure 4). K12, K155, and K446 were identified as putative ubiquitination with “medium confidence” (i.e. Score range of 0.69 to 0.84, sensitivity of 0.346, and specificity of 0.950). Two additional sites identified as “low confidence” were not indicated in Figure 4. In addition, using the GPS-SUMO webserver, four putative sumoylation sites and one sumo-interaction site were identified (Figure 4 and Table S2). We used DISULFIND tool at Machine Learning Neural Networks Group webserver (Ceroni et al. 2006) to search for possible disulfide bonding sites and did not find any sites with positive disulfide bonding state.

**Figure 4.**
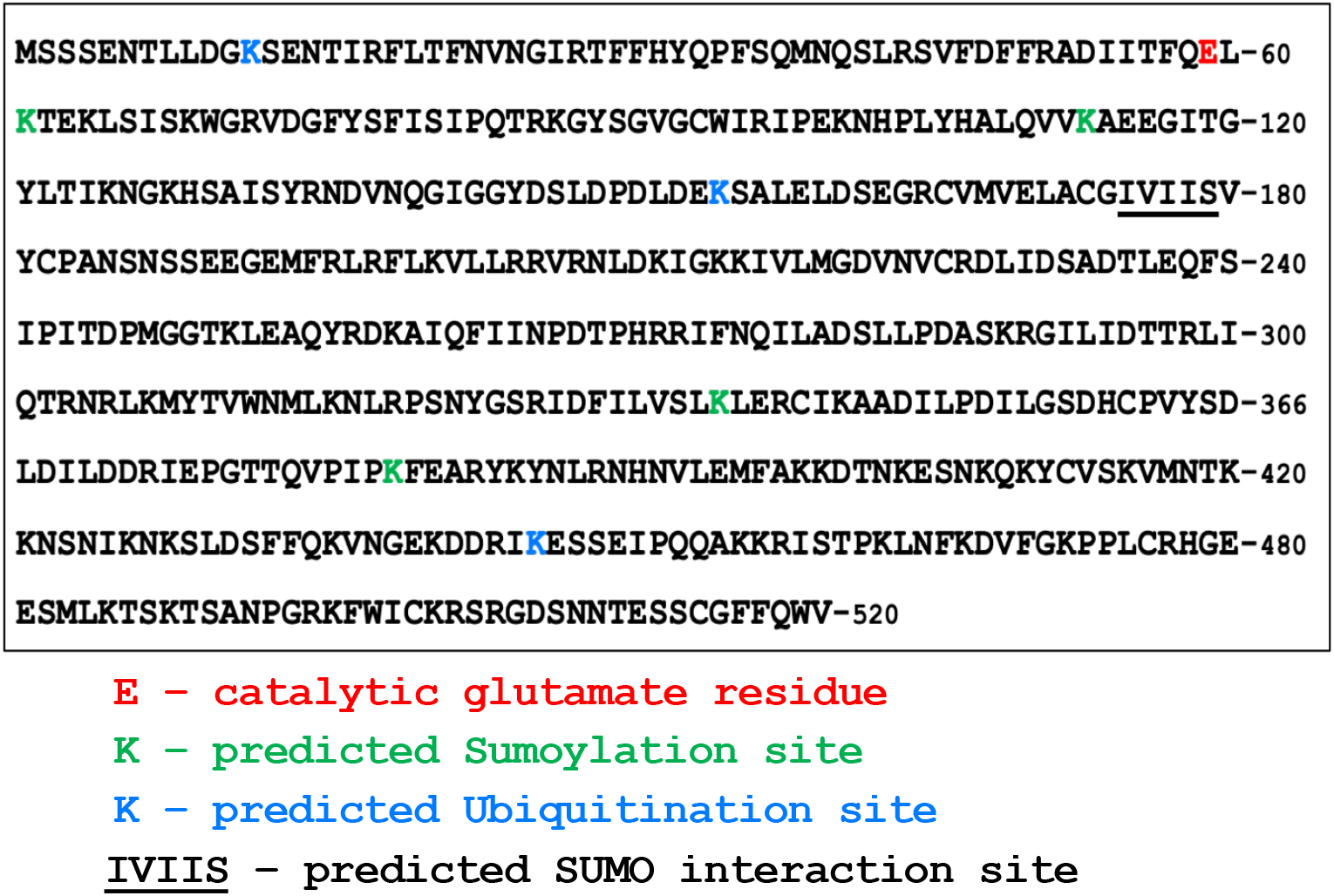
Putative Post-translational modification sites of Apn2. The protein sequence of *S. cerevisiae* Apn2 is shown in the box with the catalytic glutamate 59, putative Sumoylation sites (GPS-SUMO webserver - http://sumosp.biocuckoo.org), and putative Ubiquitination sites (UBpred - http://www.ubpred.org) indicated in red, green, and blue, respectively. Underlined residues (IVIIS) are predicted to be SUMO-interaction site (GPS-SUMO webserver - http://sumosp.biocuckoo.org).

In summary, we report here two novel findings regarding the repair of uracil-derived AP sites in the yeast genome. We show that, in absence of the major AP endonuclease Apn1, the catalytic activity of Apn2 contributes to elevation of uracil-derived A:T to C:G transversion mutations at an actively transcribed mutation reporter. Such activity of Apn2 is suppressed when yeast cells are grown in glucose, leading to the speculation that glucose repression regulates Apn2 activity. Considering the significance of uracil as one of most prevalent types of spontaneous DNA lesions, these novel findings call for future studies to determine the precise mechanism underlying such non-canonical activity of Apn2, as well as the mechanism of the glucose-dependent suppression of such activity.

## ACKNOWLEDGEMENTS

We thank the members of Kim lab for discussion and critical reading of the manuscript. This work was supported by grants from National Institutes of Health R01 GM116007 and AU1875 from Welch Foundation to N.K. A.B. was supported by R01 GM116007-S1.

