## supplemental tables 1 and 2 for "The activity of yeast Apn2 AP endonuclease at uracil-derived AP sites is dependent on the major carbon source"

Table S1. The yeast strains used in the study.

| **Strain** | **Relevant genotype** | **Source** |
|---|---|---|
| YNK98 | *apn1Δ:: loxP* | Ref. |
| YNK180 | *apn2Δ:: loxP-TRP1-loxP* | This study |
| YNK103 | *apn1Δ:: loxP apn2Δ::loxP-TRP1-loxP* | This study |
| YNK219 | *apn2Δ:: loxP-TRP1-loxP apn1Δ::loxP-URA3KL-loxP* | This study |
| YNK497 | *yap1Δ::loxP-URA3KL-loxP* | This study |
| YNK498 | *apn1Δ:: loxP yap1Δ::loxP-URA3KL-loxP* | This study |
| YNK748 | *apn1Δ:: loxP apn2Δ::loxP-Hyg*$^R$*-loxP yap1Δ::loxP-URA3KL-loxP* | This study |
| YNK500 | *dun1Δ:: loxP-TRP1-loxP* | This study |
| YNK502 | *apn1Δ:: loxP dun1Δ:: loxP-TRP1-loxP* | This study |
| YNK751 | *apn1Δ:: loxP apn2Δ::loxP-Hyg*$^R$*-loxP dun1Δ::loxP-TRP1-loxP* | This study |
| YNK787 | *rfx1Δ::loxP-Hyg*$^R$*-loxP* | This study |
| YNK581 | *apn1Δ:: loxP rfx1Δ::loxP-Hyg*$^R$*-loxP* | This study |
| YNK582 | *apn1Δ:: loxP apn2Δ::loxP-TRP1-loxP rfx1Δ::loxP-Hyg*$^R$*-loxP* | This study |
| YNK562 | *msn2Δ::loxP-Hyg*$^R$*-loxP msn4Δ::loxP-TRP1-loxP* | This study |
| YNK564 | *apn1Δ:: loxP msn2Δ::loxP-Hyg*$^R$*-loxP msn4Δ::loxP-TRP1-loxP* | This study |
| YNK790 | *apn1Δ:: loxP apn2Δ::loxP-URA3KL-loxP msn2Δ::loxP-Hyg*$^R$*-loxP msn4Δ::loxP-TRP1-loxP* | This study |
| YNK650 | *apn2 E59A-6XHA* | This study |
| YNK658 | *apn1Δ::loxP-TRP1-loxP apn2 E59A-6XHA* | This study |

All strains are derived from YNK75 [*MATα ura3-52 ade2-101oc trp1Δ1 lys2Δ::nat leu2-K:TetR'-Ssn6:LEU2 his4Δ::pTET-lys2-TAA"Same" (kan*$^R$*)*]

Table S2. The search results from GPS-SUMO webserver.

| **Type** | **Position** | **Sequence** | **Score** | **Cut-off** | **P value** |
|---|---|---|---|---|---|
| Sumoylation | 61 | QELKTEK | 12.253 | 2.13 | 0.007 |
| Sumoylation | 113 | QVVKAEE | 15.233 | 2.13 | 0.005 |
| Sumoylation | 335 | VSLKLER | 9.746 | 2.13 | 0.009 |
| Sumoylation | 379 | PIPKFEA | 6.926 | 2.13 | 0.028 |
| Sumo-interaction | 175 - 9 | ACGIVIISVY | 60.674 | 29.92 | 0.005 |
